# HiVA: an integrative wet- and dry-lab platform for haplotype and copy number analysis of single-cell genomes

**DOI:** 10.1101/564914

**Authors:** Masoud Zamani Esteki, Amin Ardeshirdavani, Daniel Alcaide, Heleen Masset, Jia Ding, Alejandro Sifrim, Parveen Kumar, Eftychia Dimitriadou, Jan Aerts, Thierry Voet, Yves Moreau, Joris Robert Vermeesch

## Abstract

Haplotyping is imperative for comprehensive analysis of genomes, imputation of genetic variants and interpretation of error-prone single-cell genomic data. Here we present a novel sequencing-based approach for whole-genome SNP typing of single cells, and determine genome-wide haplotypes, the copy number of those haplotypes as well as the parental and segregational origin of chromosomal aberrations from sequencing- and array-based SNP landscapes of single cells. The analytical workflow is made available as an interactive web application HiVA (https://hiva.esat.kuleuven.be).

---

Technologies for single-cell whole-genome analyses allow disclosing inter-cellular genetic heterogeneity^1, 2^, which is fundamentally changing our understanding of DNA mutation in development, ageing and disease^3^, and enable novel medical practice^4^, in particular for genetic selection of human preimplantation embryos^5, 6^. Current methods for single-cell genome analysis require some form of whole genome amplification (WGA) to yield sufficient input material for microarray or next-generation sequencing (NGS) analyses^1, 2, 7^. However, WGA produces artefacts –including locus drop-out (LDO), allelic drop-out (ADO), chimeric DNA molecules, base replication errors, and unevenness in amplification– that challenge the detection of genetic variation at the single-cell level^4^.

An effective way to alleviate WGA artefacts is haplotyping that connects variant alleles present on the same DNA double helix within a single cell. Several family-based^5, 8–11^ and population-based^12–16^ methods for haplotyping of single nucleotide polymorphism (SNP) genotypes^5, 8–11^ or SNP B-allele fractions (BAFs)^15, 16^ derived from bulk DNA samples have been developed. However, these methods have a number of shortcomings for single-cell SNP haplotyping. They go awry on the error-prone single-cell SNP genotypes mainly due to ADO and putative base replication errors ^8^, often ignore DNA copy number aberrations^5, 8^ and cannot distinguish mitotic from meiotic allelic imbalances. Additionally, for methods that make use of sequencing data^15^, (ultra-)deep sequencing of genomes is needed, which is cost prohibitive and computationally demanding.

Here we devised HiVA (Haplarithm inference of Variant Alleles; Fig. 1b and **Supplementary Fig. 1**) a web application for concurrent haplotyping and copy number typing of single cells^6^ in conjunction with a novel cost-effective single-cell genotyping-by-sequencing (scGBS) approach. scGBS generates reduced genomic representation libraries using a restriction enzyme (RE) that frequently cuts the amplified genome of the cell^17, 18^, followed by size selection and PCR of the shorter fragments, and finally sequencing their both ends (Fig. 1a).

**Figure 1.**
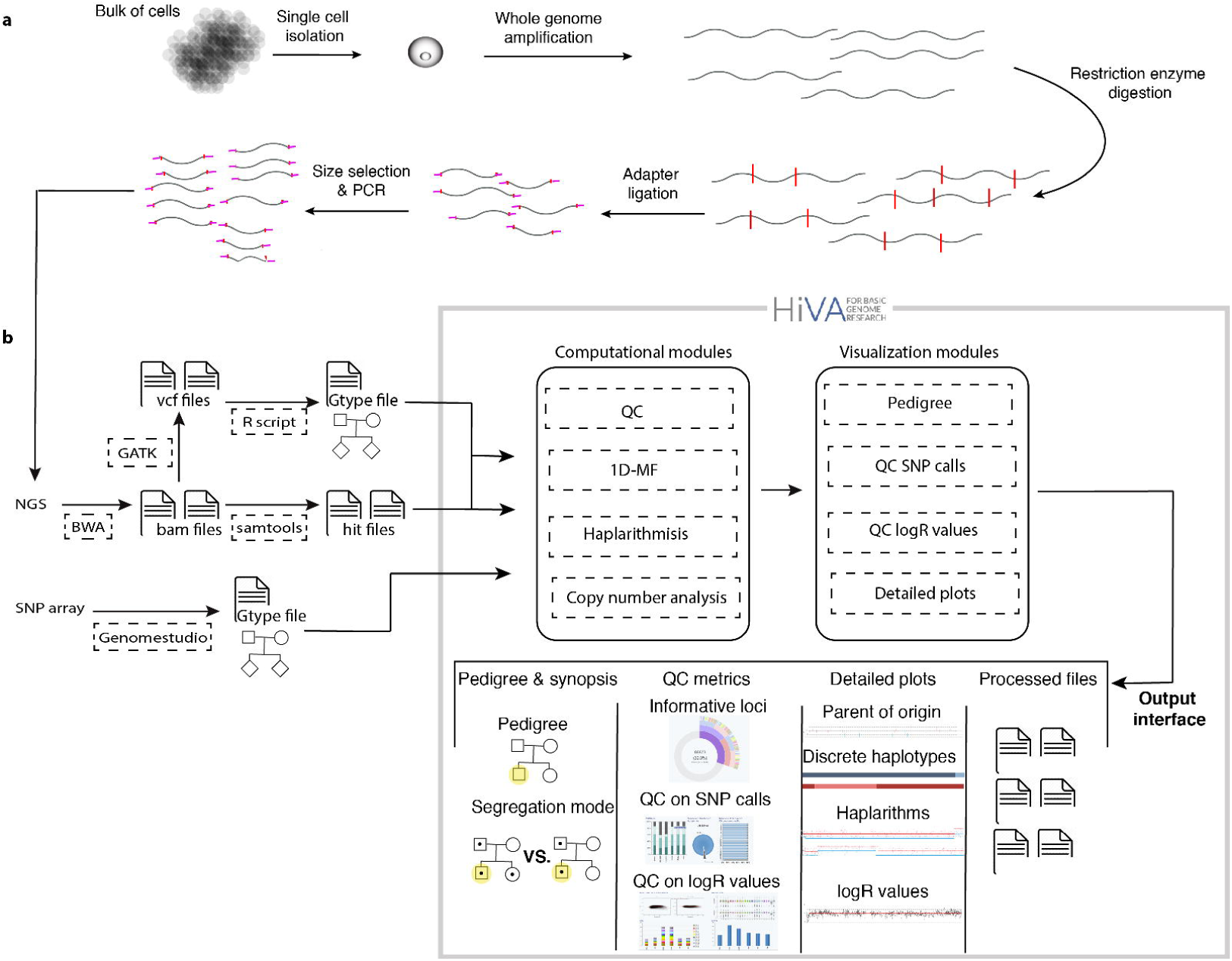
Integrative wet- and dry-lab single-cell haplotyping and copy-number profiling approach. **a**, schematic illustration of scGBS assay. **b**, overview of the interactive HiVA web application (see also **Supplementary Fig. 1**).

HiVA is developed from our previous single-cell haplotyping workflow^6^ by combining novel analytical modules, including (i) quality control filtering of both single-cell SNP-array and GBS data, (ii) family-based haplotyping of single-cell genome-wide SNP genotypes and B-allele frequencies –the latter is termed haplarithmisis^6^–, (iii) imputation of genetic variants and their distances to the nearest homologous recombination site, (iv) DNA copy number analysis of SNP-array or reduced representation genomic sequences, and (v) interactive visualization modules for integrative analysis of global or detailed single-cell haplotype-plus-copy number landscapes (Online Methods, **Supplementary Figs. 1 and 2**). As such the applications of the tool range from genome-wide haplotype reconstruction and genetic variant imputation, to deciphering the nature of allelic imbalances in single cells, including their parental origin as well as their mitotic or meiotic segregational origin^19–21^ (Fig. 2). Moreover, the methodology has been clinically implemented for genetic diagnosis of preimplantation embryos by imputation of Mendelian disease variants ^6, 21, 22^ (Fig. 2).

**Figure 2.**
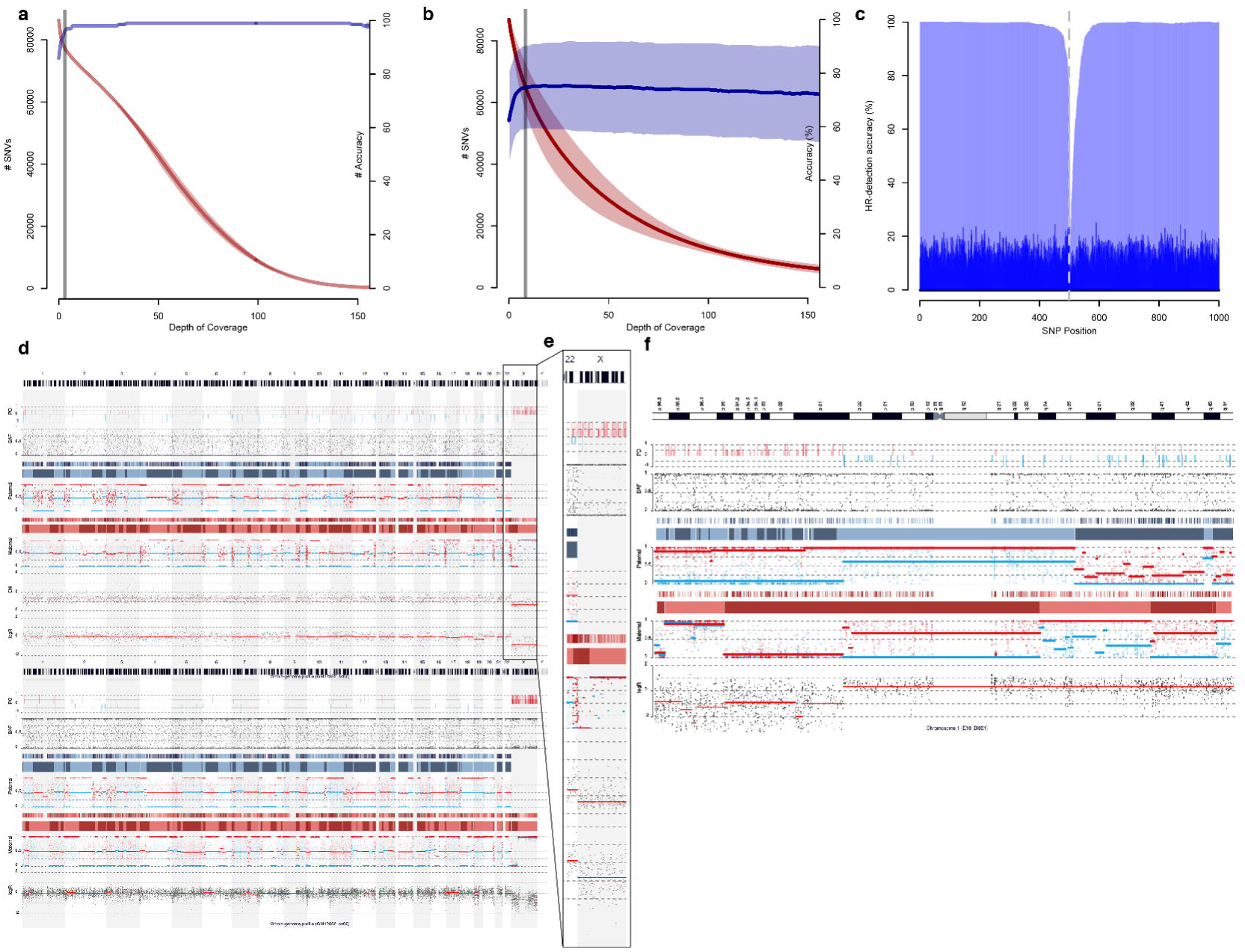
scGBS proof-of-concept assay and its application as a comprehensive PGT method. **a**, accuracy and amount of heterozygous SNP calls derived from multi-cell DNA samples when compared to their platinum SNP calls in function of depth of coverage. **b**, accuracy and amount of heterozygous SNP calls derived from single cells when compared to their multi-cell reference in function of depth of coverage. **c**, homologous recombination detection plot representing the accuracy of five single-cell haplotypes when compared to their multi-cell reference following scGBS. **d**, comparison of SNP array and scGBS haplarithms. **e**, monosomy ChrX. **f**, mitotic loss and gain of Chr1. In panels d, e, and f from top to bottom, we show chromosomes’ ideogram, parent-of-origin profile, raw BAF values, raw discrete paternal haplotypes, interpreted discrete paternal haplotypes, paternal haplarithms, raw discrete maternal haplotypes, interpreted discrete maternal haplotypes, maternal haplarithms and copy number profiles.

For performance testing of scGBS, we compared single-cell with bulk GBS sequences derived from siblings GM12882 (n=5 single cells) and GM12887 (n=2 single cells) of the CEPH/Utah 1463 HapMap family for which we also generated parental bulk GBS data. We evaluated ApeKI, NspI and PstI restriction enzymes to reduce the complexity of the whole-genome amplified single-cell genome, of which ApeKI retained the most informative SNPs (Online Methods, **Supplementary Fig. 3**, **Supplementary Note**). Following alignment of the GBS data and in silico digestion of the reference genome^23^ (Online Methods), the mean depth and breadth of sequencing coverage for the ApeKI targeted regions in the bulk DNA and single cells were 100.5 X (± 65.02 X SD) and 88.65 X (± 5.22 X SD), and 25 X (± 10.6 X SD) and 83.69 X (± 5.72 X SD), respectively (see also **Supplementary Table 1**). We first genotyped the bulk GBS sequences of individuals GM12877 and GM12878, and compared them with the available high-confidence variant calls from the Platinum genomes^24^, which showed that 7X depth of coverage produces over 98% accurate heterozygous SNP calls while retaining the majority of those SNPs (97.22% ± 0.0 SD). Using this bulk GBS as a reference, the single-cell GBS sequences produced a call rate of 73.84% (±4.11% SD) and an accuracy of 83.23% (± 7.35% SD) (Fig. 2a). Subsequently, we determined the minimal depth of coverage for scGBS by comparing its heterozygous SNP calls with the multi-cell calls in function of sequencing coverage (Fig. 2b, **Supplementary Note**), and found that 11X gives 74.71% (±15.73 SD) accurate heterozygous SNP calls while retaining the majority of those SNPs (69.67% ± 10.52 SD) (see also **Supplementary Fig. 4**). Our data showed comparable genotyping efficacy of scGBS genotype calls when compared to SNP array data of the same single-cell samples (**Supplementary Tables 2 and 3**). This single-cell and bulk SNP-information was fed to HiVA, producing single-cell haplotypes that are 99.63% (±0.63% SD) accurate when compared to bulk derived haplotypes (Fig. 2c and d). Furthermore, after transforming single-cell haplarithms to discrete haplotypes^6^, scGBS delivered comparable accuracies as compared to SNP array data (**Supplementary Tables 4 and 5**, **Supplementary Fig. 5**).

To further validate scGBS and HiVA on primary tissue, we processed 15 single blastomeres and 3 trophectoderm samples biopsied from cleavage- and blastocyst-stage human preimplantation embryos, respectively. These samples, derived from 6 families with various genetic indications (**Supplementary Table 6**), previously underwent SNP-typing with arrays for preimplantation genetic testing (PGT). All of the scGBS and SNP-array based HiVA analysis results were concordant with the clinical PGT approach (Fig. 2d and **Supplementary Table 6**).

In conclusion, here we develop a reduced representation genome sequencing approach (scGBS), enabling haplotype-specific genomic landscape profiling of single cells –without requiring full-genome deep sequencing– in combination with an innovative, user-friendly, and interactive web platform for single-cell SNP-data analysis. The methodology delivers novel understanding of chromosome instability and can potentially be translated to the clinic.

## Online content

Methods and any associated references are available in the online version of the paper.

Note: Any Supplementary Information and Source Data files are available in the online version of the paper.

## Acknowledgements

We are grateful to all families that participated in this study. We acknowledge Telerik for giving us the permission to use their interface components as well as Northwoods Software for their GoJS package. Work by A.A. and Y.M. was supported by KU Leuven: CoE PFV/10/016 SymBioSys, Flemish Government: IWT Exaptation, PhD grants, FWO 06260, and VIB: ELIXIR, imec ICON GAP, strategic funding 2017. Work by J.R.V. and T.V. is supported by KU Leuven funding (C1/018) and the Horizon 2020 WIDENLIFE: 692065 to J.R.V. and T.V. H.M. is supported by FWO (11A7119N).

## Author Contributions

M.Z.E., A.A., D.A. and devised the webtool. M.Z.E., H.M., J.D., A.A., E.D., T.V. and J.R.V. developed scGBS wet- and dry-lab protocols. A.A. and D.A. developed database management and visualization modules. M.Z.E., H.M., E.D. and J.D. collected and analyzed all the samples. P.K., A.S. and T.V. developed and implemented the focal read depth analysis pipeline. J.A., T.V., Y.M. and J.R.V. supervised the study.

## METHODS

### Definitions

Haplarithmisis (Greek for haplotype numbering) is a concept that enables haplotyping, copy number typing, parent of origin typing, and reveals segregational and mechanistic origin of genomic anomalies. n_Pat_ and n_Mat_ represent paternal and maternal copies, respectively. n_Pat_:n_Mat_ denotes the allelic ratio of a genomic region. P1 and P2 are two subcategories in the paternal haplarithm; M1 and M2 are two subcategories in the maternal haplarithm, these subcategories are determined on the basis of different parental genotype combinations ^9^. d_Pat_ represents the overall distance between P1 and P2 in a paternal haplarithm; d_Mat_ represents the overall distance between M1 and M2 in a maternal haplarithm. Haplarithmisis. Haplarithmisis uses categorization, conversion, and segmentation of BAFs from informative loci, producing P1 and P2 phased BAF segments in the paternal haplarithm (**Supplementary Fig. 2**) and similarly, M1 and M2 phased BAF segments in the maternal haplarithm. When the paternal haplotype 1 (H1) is inherited, the P1 BAF segment embodies a series of homozygous SNPs (P1 BAF = 0), while the P2 BAF segment across the same DNA locus a series of heterozygous SNPs (P2 BAF = 0.5) that are expected in the context of a diploid genome of the cell with a paternal H1 and a maternal H1 or H2 inheritance. If paternal haplotype 2 (H2) is inherited instead, the P1 BAF segment embodies a series of heterozygous SNPs (P1 = 0.5) and the P2 BAF segments a series of consecutive homozygous SNPs (P2 BAF = 1) (**Supplementary Fig. 2**). Hence, co-localizing breakpoints in the P1 and P2 segments locate homologous recombination sites from paternal meiosis, while deviations of expected P1 and P2 BAF values from an expected diploid context denote DNA copy number and ploidy anomalies (**Supplementary Fig. 2**). The vertical distance between segmented P1 and P2 in the paternal haplarithm (hd_PAT_) should complement the vertical distance between M1 and M2 in the maternal haplarithm (hd_MAT_) to the total sum amount of 1 (i.e. hd_PAT_ + hd_MAT_ = 1). These distances in combination with copy number values represent the copy number state and parental origin. For instance, for a normal diploid chromosome, hd_PAT_ and hd_MAT_ should be both 0.5. While, a chromosome that shows a copy number gain and has a hd_PAT_ value of 0.67 that is complemented with a hd_MAT_ value of 0.33 (0.67+0.33=1), represents a maternal trisomy (**Supplementary Fig. 2**).

### Human subjects

We used single-cell and multi-cell DNA samples from the lymphoblastoid cell lines derived from a HapMap Family (CEPH/Utah Pedigree 1463). We validated HiVA for single-cell analysis using single cells derived from individuals GM12882 and GM12887 of the CEPH/Utah 1463 HapMap family, as well as single-cell and few-cell biopsies from preimplantation embryos of six different PGT families.

### High-throughput single-cell genotyping-by-sequencing (scGBS)

Restriction enzyme digestion and library preparation are carried out as described previously^19^ with the following modifications to the protocol: 500ng of input DNA is used for both multi-cell DNA and single-cell WGA product, pooling occurs at equal amounts of DNA (100ng) and an additional size-selection step (140-240 bp) is performed prior to a PCR amplification of 8 cycles. Barcodes are generated through the module Barcode Generator of GBSX^12^. For *in silico* restriction enzyme digestion and subsequent determination of the amount of expected target sequence (125 nucleotides surrounding the restriction site), we applied the Restriction Enzyme Predictor tool of GBSX^12^. For HapMap samples and three PGT-M families paired-end (2×125bp) sequencing was performed on a HiSeq2500 system (Illumina) in multiple runs. For the remaining PGT-M samples, paired-end (2×150bp) sequencing was performed on a NextSeq 550 system (Illumina) with 20 samples on one run in High Output mode. Paired-end sequencing data was demultiplexed using the Demultiplexer module of GBSX^23^. Subsequently, paired-end reads were flashed (i.e. reads of fragments that in size are smaller than 2x read length that are found to be overlapping are merged) using FLASH^25^. Afterwards, both flashed and non-flashed reads could be merged and mapped to the reference genome (hg19 genome build) with BWA^26^ MEM (**Supplementary Note**). We applied GATK’s^27^ Depth of Coverage module for determining the depth across the targeted regions.

### High-throughput genotyping by SNP-array

The HumanCytoSNP-12v2.1 (Illumina; GEO: GPL13829) BeadChips were performed according to manufacturer’s instructions using genomic DNA isolated from a large number of cells. For the HapMap samples, 600 ng of single-cell single-cell WGA DNA and 200 ng multi-cell DNA isolated from a large number of cells was used. Subsequently, for genotype calling, the signal intensities were analyzed by the GenCall algorithm (http://www.illumina.com/software/genomestudio_software.ilmn) as described previously^6^.

### Genotype inference following GBS

We applied GATK’s^27^ Haplotypecaller (Best Practice) to the BAM files of each sample. For haplotype reconstruction, we applied only those SNVs having a depth of coverage of ≥ 7 for multi-cell samples and ≥ 11 for single-cell samples. A multi-sample genotyping file was created using GATK’s GVCFs. The SNV calls were then transformed to bi-allelic calls (i.e. AA, AB and, BB) with a custom R script using the following Bioconductor libraries: SNPlocs.Hsapiens.dbSNP144.GRCh37, VariantAnnotation and vcfR. BAF values were calculated based on allele-specific depth of coverage (**Supplementary Note**).

For computation of (relative) copy number values, we applied a modified version of our previous focal read-depth analysis^42^, but enabled user definable bin sizes in HiVA.

### Web interface

We used ASP.NET C# as an open source language to develop our MVC (Model-View-Control) based web application (see also **Supplementary Fig. 1**).

### Back-end application and database management

HiVA’s application server runs multiple projects concurrently. Therefore, we devised a job scheduler to make sure that all the requests are processed in an efficient way. The job controller is responsible for scheduling requests and transferring files and results between the application server, database, and web server. However, processing data by HiVA generates 28 different data files. The size of the data is thereby increased to more than two gigabytes (GB). To tackle this, we optimized a database structure to store the results and serve the web application. Our database server is a virtual machine shared on a normal server with 4 cores and 4 GB of memory, indicating the adequacy of our database structure.

### Data visualization

HiVA visualizes different QC metrics and provides genome- and chromosome-specific interactive plots with the capacity of adapting to screen sizes and adjusting the amount of visualized data points. It also provides instant access to the exact coordinates in the UCSC Genome Browser and the possibility to export the view selected by the user. Furthermore, deletions, insertions, duplications, translocations, and inversions across the genome are distinguished by employing representative glyphs and links over the chromosome ideogram. The visualization module is developed in JavaScript and renders the output directly in the HTML5 canvas element. This allows plotting hundreds of thousands of data points in a single HTML page and optimizes the performance of HiVA.

### QC metrics

HiVA provides different QC metrics on genotype SNP calls and logR values. It makes use of parental genotype calls and determines Mendelian inconsistencies, including ADO, LDO, ADI. For evaluation of noise in logR values, median absolute pairwise difference (MAPD) value is determined that measures the absolute difference of two consecutive normalized logR values^28^ as well as the cumulative standard deviation (CSD) that is summed standard deviation of each chromosome per cell.

