## Supplementary Info for "HiVA: an integrative wet- and dry-lab platform for haplotype and copy number analysis of single-cell genomes"

### **Supplementary Note**

#### **Selection of optimal restriction enzyme for scGBS assay**

Proof-of-principle experiments were performed using three different restriction enzymes (REs), including ApeKI, NspI and PstI. Each of these REs displayed a different frequency of recognition sites across the human genome (**Supplementary Fig. 3**). To evaluate the yield of informative SNPs for haplotyping a joint calling on the samples of mother and father was performed with GATK’s^1^ Haplotypecaller and the number of informative SNP calls was counted (**Supplementary Fig. 3**). Compared with the SNP array platform (Illumina Human CytoSNP12v2.1 with roughly 99K informative SNPs) used in the clinical setting^2,3^, we selected ApeKI, which is the most frequent cutter that reached the required number of informative SNP calls, to use in subsequent experiments (**Supplementary Fig. 3**).

#### **Determining optimal depth of coverage**

To measure the accuracy of genotype calls following scGBS, we first compared SNV calls of two multi-cell DNA samples with the Platinum genomes (Illumina, Inc., USA). Specifically, heterozygous Platinum SNV calls of the mother (GM12878) and the father (GM12877) of the HapMap family were compared with the corresponding genotype calls from the GBS libraries. This allowed us to determine the minimum depth of sequencing coverage to call a maximum of heterozygous SNVs reliably. This resulted in a minimum accuracy of 98% for 7X or more sequencing depth. Subsequently, we compared multi-cell heterozygous SNV GBS calls of individuals GM12882 and GM12887 (two siblings of the HapMap family) with ≥7X depth of coverage with single-cell GBS data of the same cell lines. This resulted in a minimum accuracy of 74.71% for 11X or more sequencing depth (**Fig. 2b**).

#### **scGBS data analysis**

Following scGBS of the HapMap family CEPH/Utah 1463, we demultiplexed paired-end sequencing data using GBSX Demultiplexer module^4^. Subsequently, paired-end overlapping reads were merged into one read using FLASH^5^. We then mapped merged and non-overlapping reads using BWA^6^ MEM. Samtools^7^ was applied to: (i) convert SAM to BAM files, (ii) sort the mapped reads, and (iii) create an index file. These mapped reads were merged together into one BAM file using Samtools. Default settings of GATK’s Haplotypecaller were applied to each individual sample to generate gvcf files with --emitRefConfidence set to BP_RESOLUTION. Depth filtering was applied per-sample with GATK’s VariantFiltration: for single-cell samples depth of coverage was set to ≥ 11X and multi-cell DNA samples depth ≥7X. All individual filtered samples were then genotyped together by applying GATK’s GenotypeGVCFs to generate a multi-sample gvcf file. Subsequently, we computed single-cell genotyping accuracy and WGA artefacts following both SNP-array and scGBS (**Supplementary Tables 2** and **3**). Subsequently, after transforming single-cell haplarithms, scGBS yielded comparable haplarithms as compared to SNP array (**Supplementary Tables 4** and **5**).

#### **Validation of scGBS for preimplantation genetic testing (PGT)**

In a retrospective validation study, we recruited 6 PGT families (18 embryos) and applied our integrative wet- and dry-lab approach, i.e. scGBS and HiVA. All the diagnoses were concordant with the SNP-array based approach (**Supplementary Table 6**).

### **Supplementary Tables**

#### ***Supplementary Table 1 | Amount of reads per bulk as well as few-cell and single-cell DNA samples.***

| **Sample name** | **Sample type** | **Read count** |
| --- | --- | --- |
| GM12877 | Multi cell | 16866969 |
| GM12878 | Multi cell | 16139624 |
| GM12882_MC | Multi cell | 34857794 |
| GM12882_sc37 | Single cell | 17648460 |
| GM12882_sc40 | Single cell | 15279623 |
| GM12882_sc1 | Single cell | 15089838 |
| GM12882_sc4 | Single cell | 54715575 |
| GM12882_sc6 | Single cell | 47996267 |
| GM12887_MC | Multi cell | 27340485 |
| GM12887_sc4 | Single cell | 10054700 |
| GM12882_sc6 | Single cell | 10986046 |
| PGD051_C1_E01 | Single cell | 36769818 |
| PGD051_C1_E08 | Single cell | 39929513 |
| PGD051_C1_E09 | Single cell | 42289310 |
| PGD051_F | Multi cell | 85673798 |
| PGD051_MGF | Multi cell | 34672213 |
| PGD051_MGM | Multi cell | 47061774 |
| PGD051_M | Multi cell | 106613407 |
| PGD058_C1_E01 | Single cell | 28659987 |
| PGD058_C1_E02 | Single cell | 34435992 |
| PGD058_F | Multi cell | 101110677 |
| PGD058_MGF | Multi cell | 30685496 |
| PGD058_MGM | Multi cell | 50299616 |
| PGD058_M | Multi cell | 150187874 |
| PGD088_Aff | Multi cell | 32267769 |
| PGD088_C1_E06 | Single cell | 39920083 |
| PGD088_C1_E08 | Single cell | 50155020 |
| PGD088_C1_E10 | Single cell | 46161665 |
| PGD088_C1_E13 | Single cell | 1322974 |
| PGD088_F | Multi cell | 31559241 |
| PGD088_M | Multi cell | 13600413 |
| PGD111_Aff | Multi cell | 20757315 |
| PGD111_C1_E03 | Single cell | 16514903 |
| PGD111_C1_E07 | Single cell | 13334434 |
| PGD111_C4_E03 | Single cell | 11804838 |
| PGD111_F | Multi cell | 22540339 |
| PGD111_M | Multi cell | 16955924 |
| PGD146_C3_E04 | Few cell (TE) | 17375922 |
| PGD146_C3_E08 | Few cell (TE) | 12232749 |
| PGD146_C3_E18 | Few cell (TE) | 13423582 |
| PGD146_F | Multi cell | 18252030 |
| PGD146_MGF | Multi cell | 13483230 |
| PGD146_MGM | Multi cell | 14167230 |
| PGD146_M | Multi cell | 15239966 |
| PGD230_C1_E08 | Single cell | 8665237 |
| PGD230_C1_E09 | Single cell | 11957551 |
| PGD230_C2_E02 | Single cell | 9412394 |
| PGD230_F | Multi cell | 18773035 |
| PGD230_MGF | Multi cell | 15298329 |
| PGD230_M | Multi cell | 13456392 |

TE: Trophectoderm

#### ***Supplementary Table 2 | Genotyping accuracy and WGA artifacts following SNP array***

| **Single cell** | **CC** | **ADO** | **ADI** | **ADOI** | **LDO** | **CNM** | **Acc.** |
| --- | --- | --- | --- | --- | --- | --- | --- |
| **GM12882_sc01** | 53.203 | 8.752 | 0.103 | 0.012 | 37.819 | 0.11 | 85.715 |
| **GM12882_sc04** | 64.228 | 3.689 | 0.028 | 0.002 | 31.939 | 0.114 | 94.528 |
| **GM12882_sc06** | 69.586 | 0.518 | 0.045 | 0.001 | 29.64 | 0.156 | 99.197 |
| **GM12882_sc37** | 62.169 | 3.702 | 0.042 | 0.002 | 33.928 | 0.158 | 94.317 |
| **GM12882_sc40** | 50.445 | 8.129 | 0.095 | 0.002 | 41.192 | 0.136 | 85.978 |
| **GM12887_sc04** | 56.979 | 6.428 | 0.087 | 0.001 | 36.355 | 0.15 | 89.738 |
| **GM12887_sc06** | 51.774 | 8.303 | 0.159 | 0.002 | 39.614 | 0.147 | 85.948 |

ADO: Allelic drop out; ADI: Allelic drop in; ADOI: allelic dropout followed by allelic drop in

LDO: locus drop out (dropout of both alleles) / no calls

CC: Correctly called SNP calls from all the SNP probes in the single cells when compared to their multi-cell reference

Acc: Correctly called SNP calls from called SNP probes in the single cells when compared to their multi-cell reference

CNM: called in single cell but not in bulk

#### ***Supplementary Table 3 | Genotyping accuracy and WGA artifacts following scGBS***

| **Single cell** | **CC** | **ADO** | **ADI** | **ADOI** | **LDO** | **CNM** | **Acc.** |
| --- | --- | --- | --- | --- | --- | --- | --- |
| **GM12882_sc01** | 57.316 | 10.24 | 5.415 | 0.607 | 24.532 | 1.89 | 77.899 |
| **GM12882_sc04** | 72.739 | 2.593 | 1.753 | 0.202 | 20.867 | 1.846 | 94.115 |
| **GM12882_sc06** | 70.612 | 3.532 | 2.053 | 0.232 | 21.76 | 1.81 | 92.388 |
| **GM12882_sc37** | 57.76 | 7.744 | 3.11 | 0.243 | 29.612 | 1.531 | 83.884 |
| **GM12882_sc40** | 49.642 | 13.448 | 2.527 | 0.539 | 32.303 | 1.541 | 75.037 |
| **GM12887_sc04** | 56.908 | 11.376 | 2.375 | 0.416 | 27.341 | 1.584 | 80.067 |
| **GM12887_sc06** | 56.808 | 12.142 | 2.405 | 0.368 | 26.685 | 1.593 | 79.206 |

ADO: Allelic drop out; ADI: Allelic drop in; ADOI: allelic dropout followed by allelic drop in

LDO: locus drop out (dropout of both alleles) / no calls

CC: Correctly called SNP calls from all the SNPs in the single cells when compared to their multi-cell reference

Acc.: Correctly called SNP calls from called SNPs in the single cells when compared to their multi-cell reference

CNM: called in single cell but not in bulk

#### ***Supplementary Table 4 | Haplarithm accuracy following SNP array.***

|  |  | Paternal | | | Maternal | | |
| --- | --- | --- | --- | --- | --- | --- | --- |
| Single cell | Total SNPs  (#) | Deduced haplotypes (matched P1 and P2)  (#) | Deduced haplotypes (matched P1 and P2)  (%) | Accuracy  (%) | Deduced haplotypes (matched M1 and M2)  (#) | Deduced haplotypes (matched M1 and M2)  (%) | Accuracy  (%) |
| **GM12882_sc01*** | 98636 | 87763 | 87.04 | 99.98 | 88770 | 87.64 | 99.80 |
| **GM12882_sc04** | 98636 | 92991 | 92.20 | 100.00 | 95729 | 94.53 | 100.00 |
| **GM12882_sc06** | 98636 | 93827 | 92.93 | 100.00 | 96356 | 95.19 | 100.00 |
| **GM12882_sc37** | 98636 | 92715 | 92.16 | 100.00 | 94777 | 94.11 | 100.00 |
| **GM12882_sc40*** | 98636 | 91348 | 90.94 | 99.98 | 93723 | 93.03 | 100.00 |
| **GM12887_sc04** | 98688 | 92227 | 92.11 | 100.00 | 94761 | 94.07 | 100.00 |
| **GM12887_sc06** | 98688 | 91783 | 91.69 | 99.97 | 94534 | 93.73 | 100.00 |

*Samples that did not pass CSD QC metric of HiVA.

#### ***Supplementary Table 5 | Haplarithm accuracy following scGBS.***

|  |  | Paternal | | | Maternal | | |
| --- | --- | --- | --- | --- | --- | --- | --- |
| Single cell | Total SNPs  (#) | Deduced haplotypes (matched P1 and P2)  (#) | Deduced haplotypes (matched P1 and P2)  (%) | Accuracy  (%) | Deduced haplotypes (matched M1 and M2)  (#) | Deduced haplotypes (matched M1 and M2)  (%) | Accuracy  (%) |
| **GM12882_sc01*** | 96730 | 46089 | 46.20 | 98.73 | 49137 | 49.54 | 97.93 |
| **GM12882_sc04** | 96730 | 86663 | 87.06 | 99.91 | 85821 | 87.04 | 99.80 |
| **GM12882_sc06** | 96730 | 89584 | 90.61 | 100.00 | 91106 | 93.01 | 99.92 |
| **GM12882_sc37** | 96730 | 89559 | 89.95 | 100.00 | 88799 | 90.47 | 99.80 |
| **GM12882_sc40*** | 96730 | 90008 | 90.86 | 90.86 | 91183 | 93.21 | 99.98 |
| **GM12887_sc04** | 97064 | 87850 | 89.67 | 99.95 | 88361 | 90.21 | 99.9 |
| **GM12887_sc06** | 97064 | 88247 | 90.15 | 99.85 | 88642 | 90.59 | 99.92 |

*Samples that did not pass CSD QC metric of HiVA.

#### ***Supplementary Table 6 | PGT families involved in this study***

| **PGT family** | **PGT ID** | **Indication (gene)** | **Inheritance mode** | **Family member** | **Disease status** |
| --- | --- | --- | --- | --- | --- |
| 1 | PGD051-M | BRCA2 | AD | Mother | affected |
|  | PGD051-F |  |  | Father | normal |
|  | PGD051-MGM |  |  | Maternal Grandmother | affected |
|  | PGD051-MGF |  |  | Maternal Grandfather | normal |
|  | PGD051_C1_E01 |  |  | Embryo | unaffected |
|  | PGD051_C1_E08 |  |  | Embryo | affected |
|  | PGD051_C1_E09 |  |  | Embryo | affected |
| 2 | PGD088-M | CFTR | AR | Mother | carrier |
|  | PGD088-F |  |  | Father | carrier |
|  | PGD088-Aff |  |  | Sibling | affected |
|  | PGD088_C1_E06 |  |  | Embryo | carrier |
|  | PGD088_C1_E08 |  |  | Embryo | Inconclusive (HR)* |
|  | PGD088_C1_E10 |  |  | Embryo | carrier |
|  | PGD088_C1_E13 |  |  | Embryo | affected |
| 3 | PGD058-M | RP3 | XLR | Mother | carrier |
|  | PGD058-F |  |  | Father | normal |
|  | PGD058-MGM |  |  | Maternal Grandmother | carrier |
|  | PGD058-MGF |  |  | Maternal Grandfather | normal |
|  | PGD058_C1_E01 |  |  | Embryo | carrier female |
|  | PGD058_C1_E02 |  |  | Embryo | affected male |
| 4 | PGD111_M | DMPK | AD | Mother | affected |
|  | PGD111_F |  |  | Father | normal |
|  | PGD111_Aff |  |  | Sibling | affected |
|  | PGD111_C1_E03 |  |  | Embryo | affected |
|  | PGD111_C1_E07 |  |  | Embryo | unaffected |
|  | PGD111_C4_E03 |  |  | Embryo | inconclusive (HR)^*^ |
| 5 | PGD146_M | SCN5A | AD | Mother | affected |
|  | PGD146_F |  |  | Father | normal |
|  | PGD146_MGM |  |  | Maternal Grandmother | normal |
|  | PGD146_MGF |  |  | Maternal Grandfather | affected |
|  | PGD146_C3_E04 |  |  | Embryo | unaffected |
|  | PGD146_C3_E08 |  |  | Embryo | affected |
|  | PGD146_C3_E18 |  |  | Embryo | unaffected |
| 6 | PGD230_M | PKD1 | AD | Mother | affected |
|  | PGD230_F |  |  | Father | normal |
|  | PGD230_MGM |  |  | Maternal Grandmother | affected |
|  | PGD230_MGF |  |  | Maternal Grandfather | normal |
|  | PGD230_C1_E08 |  |  | Embryo | affected |
|  | PGD230_C1_E09 |  |  | Embryo | unaffected |
|  | PGD230_C2_E02 |  |  | Embryo | unaffected |
| AD: Autosomal Dominant; AR: Autosomal Recessive; XLR: X-linked recessive; Bl: Blastomere; | | | | | |
| *: inconclusive result due to the presence of a homologous recombination site (HR) too close to the locus of interest | | | | | |

### **Supplementary Figures**

#### **Supplementary Figure 1 | HiVA’s architecture.**

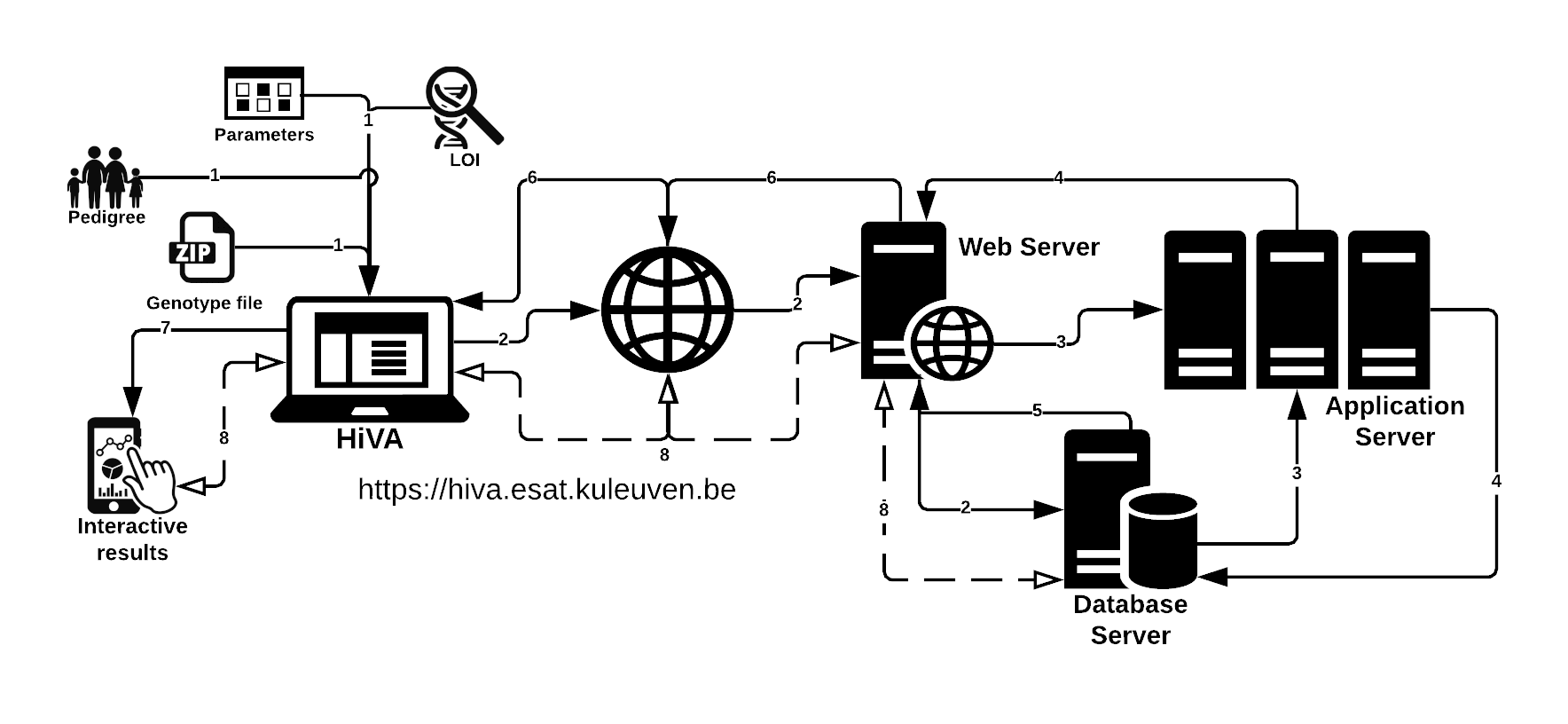

#### ***Supplementary Figure 2 | Principles of haplarithmisis process.***

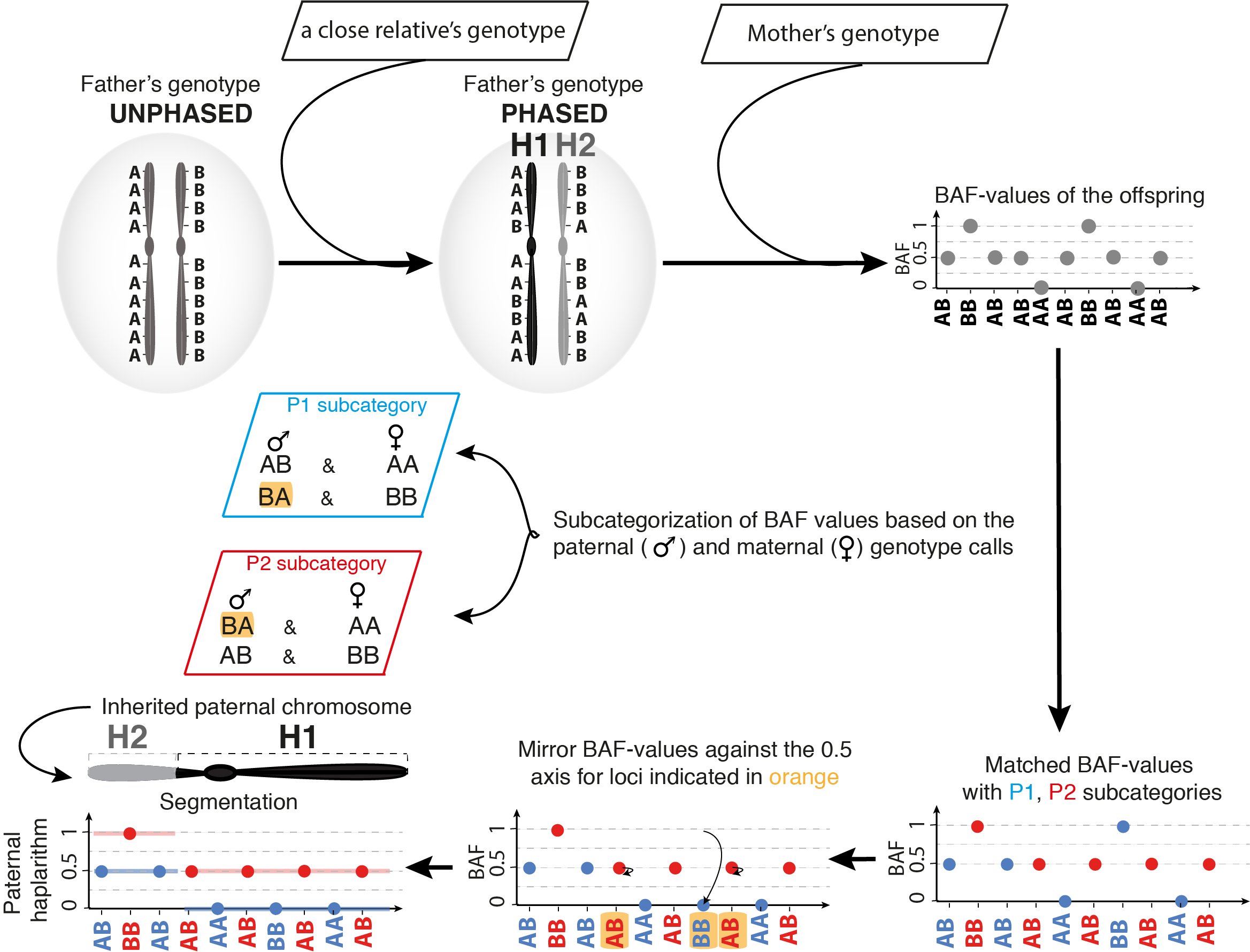

**Supplementary Figure 3 | Fragment size distribution of the three different restriction enzymes.** Fragment size profiles generated for each of the three restriction enzymes, ApeKI, NspI and PstI, tested in the proof-of-principle experiments for the scGBS assay after *in silico* digestion with GBSX’s^4^ Restriction Enzyme Predictor module. Dependent on the frequency of recognition sites for each of the REs, a different number of fragments per size category is generated. ApeKI is the most frequent cutter of the REs, thereby generating mostly smaller fragments. Relatively, NspI is an intermediate cutter and PstI is the least frequent cutter, generating a high percentage of fragments >= 1000 base pairs.

Fragment size distribution in silico digestion

Percentage

|  | ApeKI | NspI | PstI |
| --- | --- | --- | --- |
| 0-99 | 23.7518 | 12.9448 | 7.50921 |
| 100-199 | 15.0417 | 9.17104 | 5.68002 |
| 200-299 | 10.6883 | 8.64309 | 5.00125 |
| 300-399 | 8.01846 | 7.23588 | 4.55491 |
| 400-499 | 6.07674 | 6.11808 | 4.02423 |
| 500-599 | 4.95493 | 5.41479 | 3.68523 |
| 600-699 | 4.16827 | 4.9228 | 3.80839 |
| 700-799 | 3.46572 | 4.4045 | 3.27485 |
| 800-899 | 3.04353 | 3.95079 | 2.95445 |
| 900-999 | 2.55215 | 3.53251 | 2.87914 |
| >=1000 | 18.2384 | 33.6617 | 56.6283 |

#### **Supplementary Figure 4 | Targeted breadth-of-coverage (%) at different sequencing depth-of-coverage (X).** Red indicates multi-cell DNA sample and Blue indicates single-cell DNA samples (error bars represent standard deviation).

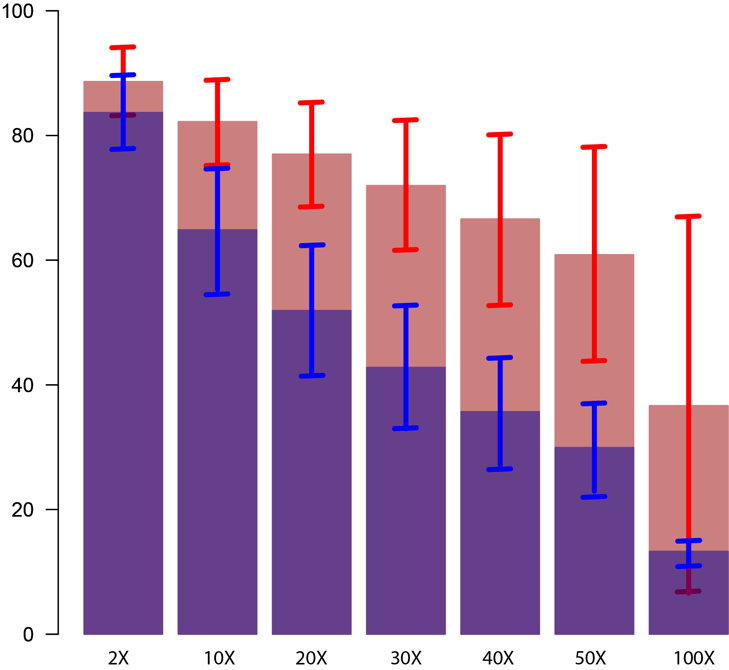

#### **Supplementary Figure 5 | SNP array versus scGBS.** HiVA plots following SNP-array (top panel) and scGBS (bottom panel) analysis of the single cell GM12882_sc06.

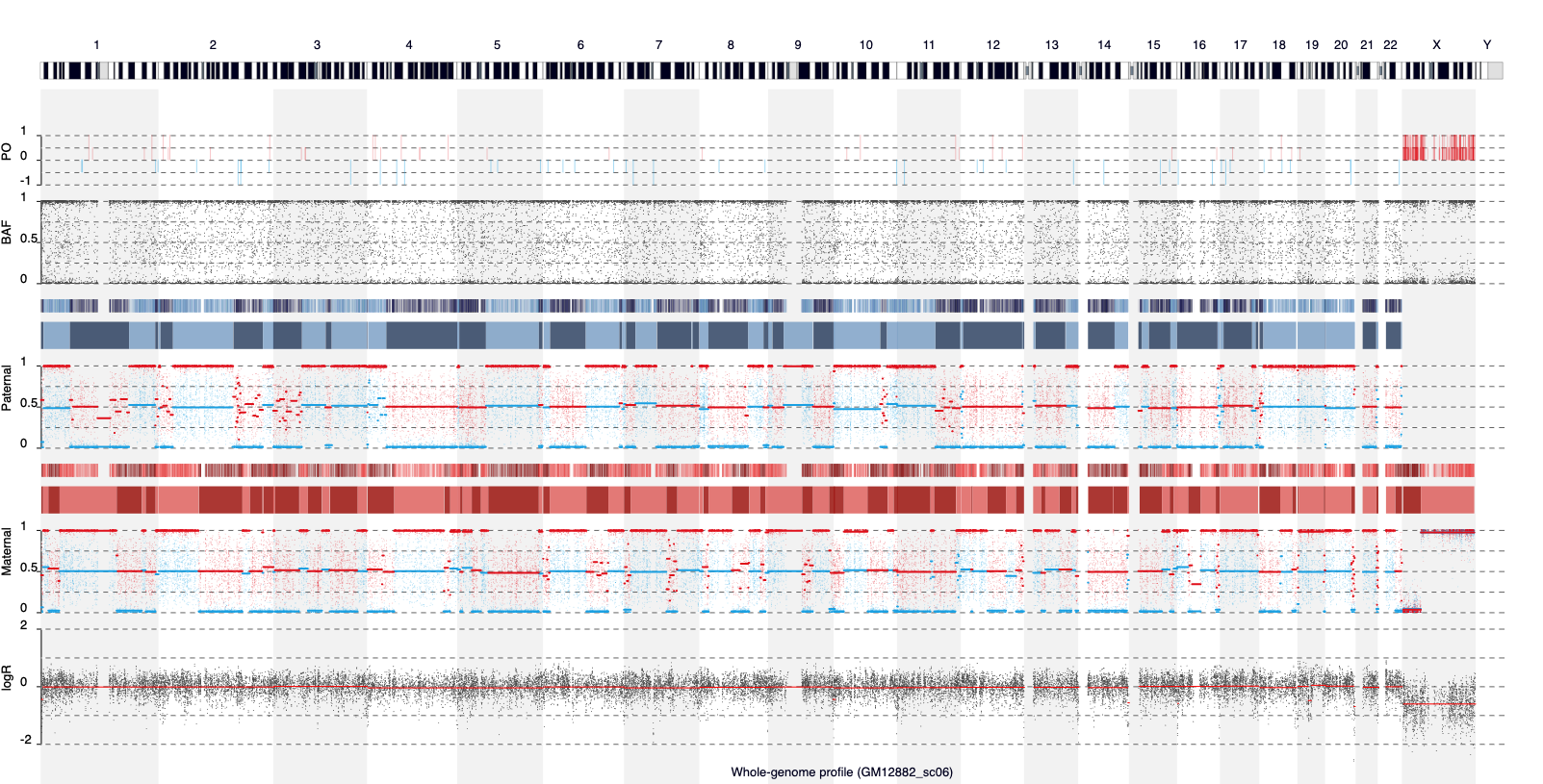

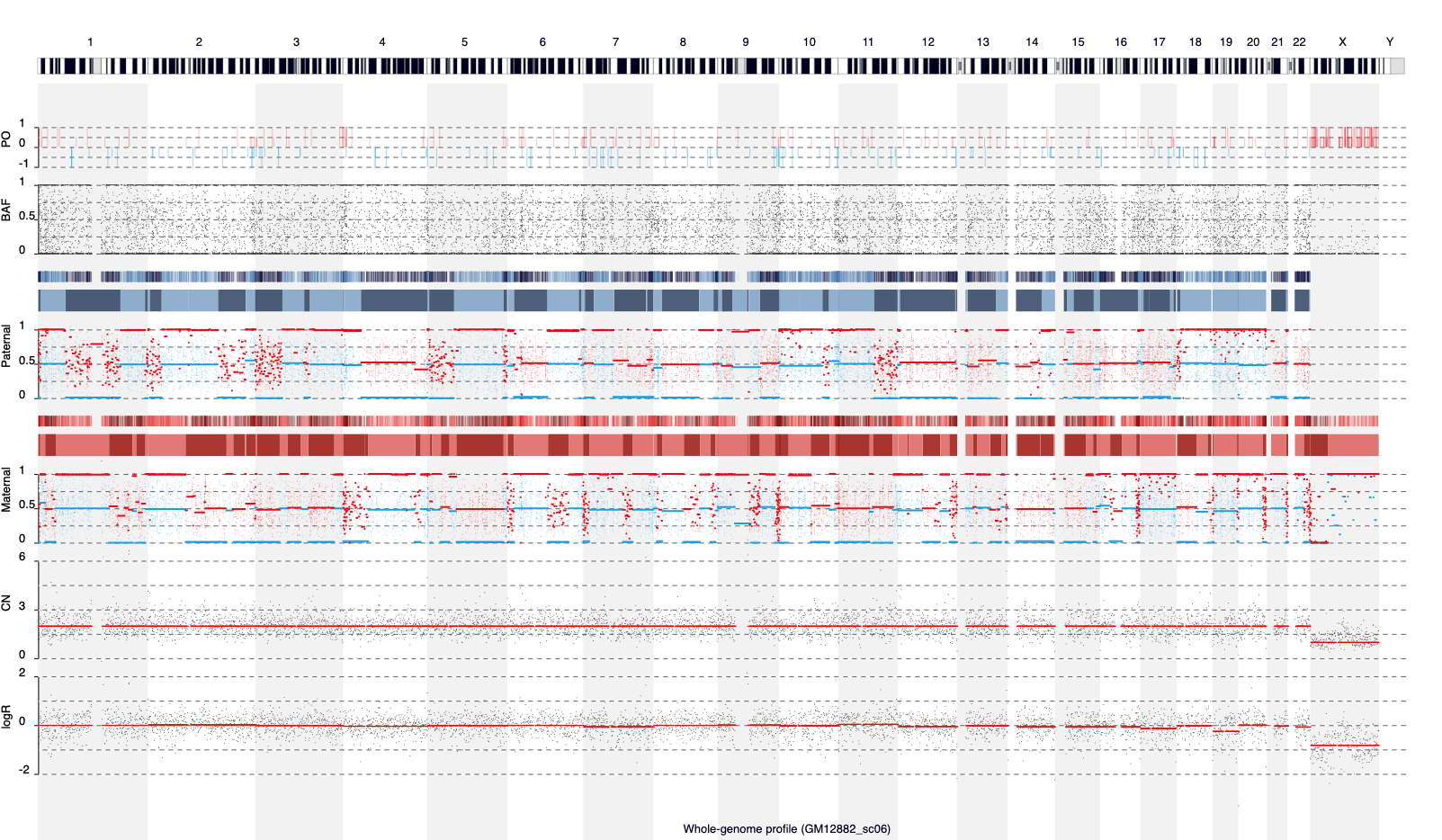
